# Disinfection performance of a drinking water bottle system with a UVC LED cap against waterborne pathogens and heterotrophic contaminants

**DOI:** 10.1101/2021.06.02.446792

**Authors:** Richard M. Mariita, Sébastien A. Blumenstein, Christian M. Beckert, Thomas Gombas, Rajul V. Randive

## Abstract

**Background:** The purgaty One systems (cap+bottle) are portable stainless-steel water bottles with ultraviolet subtype C (UVC) disinfection capability. This study examines the bottle design, verifies disinfection performance against *Escherichia coli*, *Pseudomonas aeruginosa*, *Vibrio cholerae* and heterotrophic contaminants and addresses the public health relevance of heterotrophic bacteria.

**Methods:** Bottles were inoculated with bacterial strains and disinfection efficacy examined using colony forming unit (CFU) assay. The heterotrophic plate count (HPC) method was used to determine the disinfection performance against environmental contaminants at day 0 and after 3 days of water stagnation. All UVC irradiation experiments were performed under stagnant conditions to confirm that the preset application cycle of 55 seconds offers the desired disinfection performance under worst case condition. To determine effectiveness of purgaty One systems (cap+bottle) in disinfection, inactivation efficacy or log reduction value (LRV) was determined using bacteria concentration between UVC ON condition and controls (UVC OFF). The study utilized the 16S rRNA gene for isolate characterization by identifying HPC bacteria to confirm if they belong to groups that are of public health concern.

**Results:** Purgaty One systems fitted with Klaran UVC LEDs achieved 99.99% inactivation (LRV4) efficacy against *E. coli* and 99.9% inactivation (LRV3) against *P. aeruginosa*, *V. cholerae* and heterotrophic contaminants. Based on the 16S rRNA gene analyses, the study determined that the identified HPC isolates enriched by UVC irradiation are of rare public health concern.

**Conclusion:** The bottles satisfactorily inactivated the target pathogenic bacteria and HPC contaminants even after 3 days of water stagnation.

## 1. Introduction

The low quality of potable water is a major issue in travel medicine, especially when visiting places with poor hygienic conditions due to waterborne diseases which pose substantial health risk [1]. Waterborne pathogens, predominantly of fecal origin, can be transmitted via contaminated drinking water [2]. Even in developed countries, they represent a risk to recreational travelers who have to rely on surface water [1]. For instance, in the United States, it is estimated that each year 560,000 people suffer from severe waterborne diseases due to the consumption of contaminated drinking water, with 7.1 million suffering from mild to moderate infections, resulting in estimated 12,000 deaths a year [3]. Hikers and campers are also exposed to waterborne disease risks if they consume untreated water from rivers and lakes [4]. Diarrheal infections are a major inconvenience in the wilderness during hiking or camping and can easily spread via contaminated water supplies and from person-to-person.

One way to prevent waterborne diseases for healthy travelling in regions with unsafe or underdeveloped water sources, is by ensuring adequate supply of potable water. Alternatively, outdoor enthusiasts can use portable and germicidal devices that ensure inactivation of microbial contaminants. Ultraviolet irradiation in the UVC range (200-280nm) has demonstrated effective inactivation of microbial contaminants in water [5]. Specifically, as part of the effort to accelerate the sustainable development goals (SDG) such as clean water and sanitation goal (SDG #6) [6], regulating microbial load is required to control waterborne diseases caused by microorganisms such as *Pseudomonas aeruginosa*, *Escherichia coli* and *Vibrio* spp. [7].

Cholera, caused by *Vibrio cholerae* remains a serious risk in emerging economies where sanitation is poor, health care limited, and drinking water unsafe [8]. Additionally, due to global warming, there is association between the spread of pathogenic vibrios and emergence of human diseases towards the temperate world [9]. Existing water infrastructure including electronic faucets can act as reservoirs and sources of outbreaks once contaminated. Hospital water, for instance, can disseminate opportunistic pathogens such as *Pseudomonas aeruginosa* and fecal coliforms, where *Escherichia coli* is a key species [10].

Furthermore, heterotrophic bacteria are a concern in drinking water systems if the counts are consistently > 500 CFU/mL. They can be an indication of general decrease in water quality and potential biofilm formation in municipal water [11]. Therefore, to eradicate elevated levels of HPC and other pathogens, portable bottle devices with disinfection features can act as a form of disinfectant.

With more than 1 billion people globally having no access to potable water, and 2.4 billion people still living in areas without adequate sanitation systems [12], there is need for portable, durable, appealing, personal and highly germicidal devices to help curb enteric pathogens. Use of UVC radiation is one of the disinfection methods recognized by WHO [12]. Unlike most methods, UV disinfects by striking the target microorganism with sufficient dose of energy, while neither altering the water, nor providing any residue [12]. UV is subdivided into three distinct bands: UVA with a wavelength of 315-400 nm, UVB with 280-315 nm, and UVC with 100-280 nm [13]. The UVC region has been found to be effective against waterborne pathogens [14]. Specifically, the wavelength range between 250-270nm, is strongly absorbed by the nucleic acids of microbial cells [15]. Theoretically, the disinfection performance of a UVC device is a function of the intensity of UVC light (irradiance) and time of exposure resulting in a UVC dose. Greater disinfection efficacy is expected at higher UVC dose [16].

The purpose of this study was to investigate the disinfection performance of the recent commercial development of the portable purgaty^®^ One system (cap+bottle) by analyzing test bottles against pathogens and heterotrophic contaminants. This study was carried out using US municipal drinking water supplied by Cohoes Water Department in New York State.

## 2. Materials and Methods

There are two bottle types of purgaty^®^ One on the market, a 650 mL and a 500 mL version. The purgaty^®^ brain (cap) can fit either of the bottles. The cap is rechargeable (Fig. 1 [b]).

**Fig.1:**
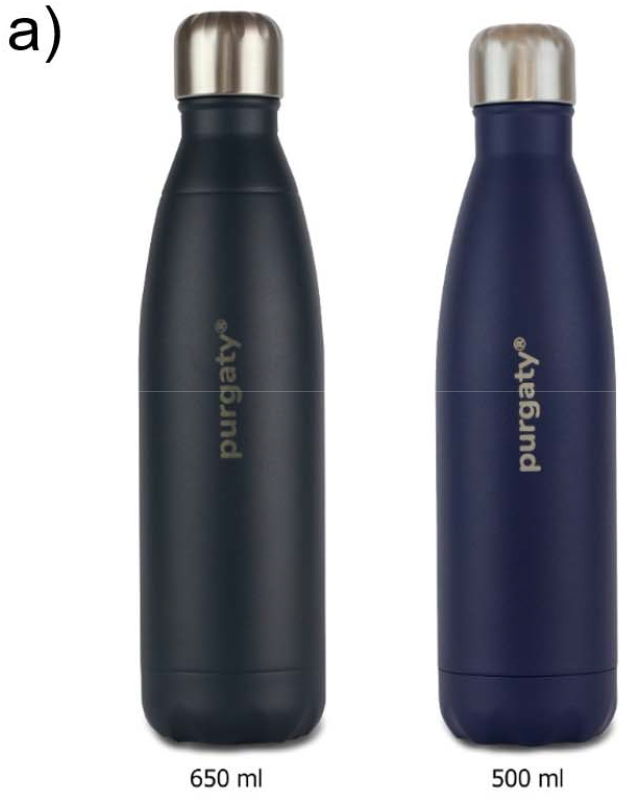
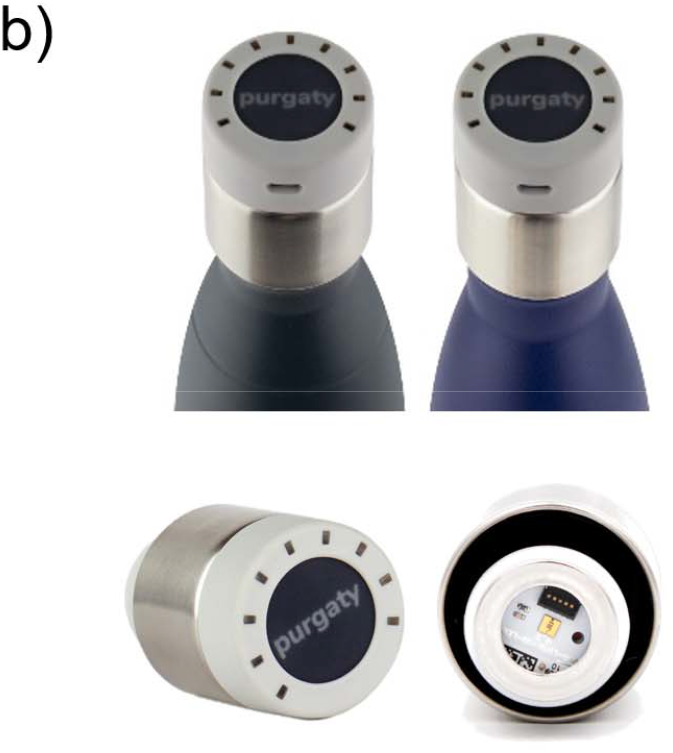
**[a].** Test bottles whose ordinary caps can be replaced with UVC emitting purgaty^®^ brain [**b].** The UVC emitting purgaty^®^ brain threaded onto both bottles (top) and shown individually from a side view and from the bottom (bottom)

### 2.1 Disinfection Cap (purgaty^®^ brain) Design

The stainless-steel bottles (Fig. 1 [a]) fitted with the UVC emitting purgaty^®^ brain (Fig. 1 [b]) offered continuous disinfection for 55 seconds. The purgaty^®^ brain is operated by means of a button on the top of the housing (Fig. 1 [b] and **Fig. 2 [a]**). Pressing the button once puts the device in standby mode, while the current charge level of the integrated battery is displayed. A second press of the button for 2 seconds activates the preset disinfection cycle for duration of 55 seconds. The unit is equipped with a safety feature, which allows the disinfection cycle to be started only after the cap has been correctly placed on the bottle (Fig. 2 [a]). A light sensor on the Printed Circuit Board (PCB) next to the UVC LED detects ambient light and interrupts the activated UVC LED in case of unthreading the cap or system damage with light entrance during a running disinfection cycle. This mechanism is designed to protect the user from contact with UVC radiation on skin or eyes. During the cycle, the LED and the sensor are periodically monitored to prevent malfunction. At the end of the 55 second cycle, the purgaty^®^ brain flashes to indicate the end of the treatment process. The cap has one Klaran UVC LED (part number KL265-50U-SM-WD) that emits at 268.5 nm peak wavelength (Fig. 2 [b]) as confirmed using an Ocean Optics USB4000 photospectrometer.

**Fig. 2:**
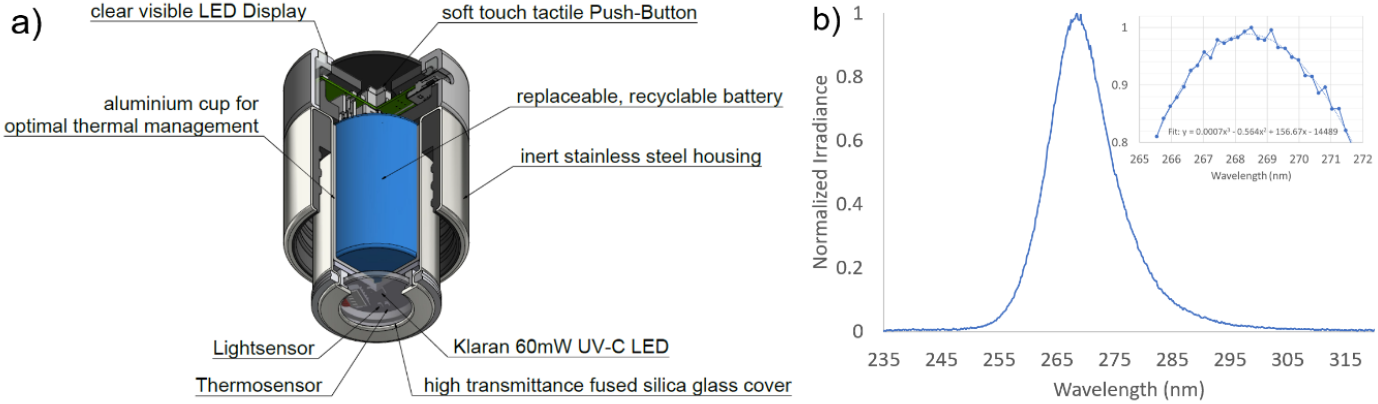
[**a]** The purgaty^®^ brain (cap emitting UVC) schematic **[b]** Measured optical spectrum of the UVC LED emission from the cap. Inset: Zoom into the peak wavelength range. A trend line has been added to the data to determine the peak wavelength value.

The thermal design is crucial for lifetime management (irreversible power degradation) as well as for ensuring the effective optical output power level (reversible thermal derating) of the UVC LED for achieving the target disinfection performance. In the purgaty^®^ brain, an aluminum cup is used to transfer the heat from the PCB to the outer stainless-steel shell, which releases the heat to the ambient air. This design ensures that after a continuous disinfection cycle of 55 seconds, the temperature on the aluminum PCB on which the UVC LED is mounted, does not exceed 45°C at an ambient temperature of 25°C maintaining enough optical output power levels for reliable disinfection performances. A plan-parallel 2 mm thick fused silica window covers the UVC LED and seals the electronics section from water and humidity. The LED package is located at 2 mm distance from the window.

### 2.2 Bacterial cultivation and enumeration of microorganism

Three strains of *Escherichia coli* ATCC 8739, *Pseudomonas aeruginosa* ATCC 15442, *Vibrio cholerae* ATCC 25872 were obtained from ATCC (Manassas, VA, USA). Stock cultures for *E. coli* and *V. cholerae* were propagated in ATCC Medium 3: Nutrient agar or nutrient broth. *P. aeruginosa* was propagated in ATCC Medium 18: Trypticase Soy Agar/Broth. All strains were plated and incubated at 37°C for 24 h. One isolated colony was picked using sterile inoculation loop and used to inoculate 25 mL broth. Flasks wi**th** side baffles were used to enhance aeration. Cultures were incubated for 18-20 h at 37°C while shaking at 180 rpm. Culture storage was done at −80°C (0.7 mL of culture with 0.7 mL of sterile 40% Glycerol stock). To obtain working cultures, the microorganisms were obtained from −80°C, streaked onto corresponding agar and incubated under same conditions. Storage of test cultures was done at −4°C. For UVC disinfection experimental use, each strain was harvested by centrifugation at 4000 rpm for 10 min. The pellets were washed using 1X PBS three times for 10 min.

Between each wash, the supernatant was discarded, and the remaining pellet re-suspended by vortexing. After washing thrice, the pellet was resuspended in 1X PBS and used to spike dechlorinated test water to achieve a final UVT value of 96% (contaminated drinking water). Dechlorination of test water was verified with a Hach DPD Free Chlorine colorimetric test. All the other test water characteristics were within the NSF/ANSI 55 standard (Tables S1-S3). The standard covers UVC disinfection systems within the range of 240 nm and 300 nm for point-of-use (POU) and point-of-entry (POE) applications [17].

### 2.3 Disinfection Experiments

The inactivation efficiency of the purgaty^®^ One system was evaluated by inoculating the bottles with 650 mL or 500 mL of contaminated water with a UVT of 96%. A single preset disinfection cycle of 55 Seconds was applied, following the manufacturer’s instructions on how to use the tactile push-button of the purgaty^®^ brain. Positive control bottles had no UVC activated as there was no use of the push-button, whereas negative control bottles contained uninoculated potable test water. The samples were thoroughly mixed after disinfection, serially diluted and processed for plating.

### 2.4 Decontamination of Heterotrophic Plate Count (HPC) bacteria

The Heterotrophic plate count (HPC) is an analytic method used to measure the variety of bacteria commonly found in water. It has no health effects as the lower the concentration of bacteria in drinking water, the better maintained the water system is. Experiments on inactivation of heterotrophic contaminants using purgaty^®^ One system were conducted on day 0 (no water stagnation in bottles), and after 3 days of water stagnation in bottles. The standard HPC technique was used and incubation at 22°C and 37°C was applied using R2A agar [18].

#### 2.4.1 HPC Isolation, Identification and Phylogenetic Analyses

Characteristic colonies (Table 2) were picked from the R2A agar plates and re-streaked for purity, incubated at 22°C prior to being shipped for sequencing. Submitted colony samples underwent a crude Sodium hydroxide lysis and were directly used in PCR. PCR amplification was performed according to Genewiz proprietary protocol. Following amplification, enzymatic cleanup was performed prior to primer extension sequencing (GENEWIZ, Inc., South Plainfield, NJ, USA) using the Applied Biosystems BigDye version 3.1. The reactions were then run on an Applied Biosystem’s 3730xl DNA Analyzer. The primer set used in this study amplifies regions V1-V9 of the 16S gene which is roughly a 1400 base pairs amplicon. Internal sequencing primers were utilized in order to allow for the generation of a consensus sequence with the forward and reverse traces. Consensus files were quality trimmed to remove the N’s. The generated 16S rRNA gene sequences were then compared with those obtained from the NCBI database, using the program BLASTN 2.2.27 + (https://blast.ncbi.nlm.nih.gov/Blast.cgi). For phylogenetic analysis, multiple alignment of *Acinetobacter* 16S rRNA gene sequences using ClustalW algorithm and tree was constructed using MEGA-X [19]. The sequences from type strains used for phylogenetics were retrieved from GenBank (National Centre for Biotechnology Information; http://www.ncbi.nlm.nih.gov/), except for the 16S rRNA gene sequences obtained through this study. *Psychrobacter cryohalolentis* K5 (Accession # NR_075055.1) was used for rooting.

## 3. Results

### 3.1 Disinfection Performances Against *Escherichia coli* ATCC 8739, *Pseudomonas aeruginosa* ATCC 15442 and *Vibrio cholerae* ATCC 25872

Table 1 shows the purgaty^®^ One system effectiveness against test strains for both bottle volume types. In all cases, both test bottles obtained a LRV greater than 3 (equivalent to greater 99.9% reduction) against target microbes. Additionally, the study revealed that *E. coli* ATCC 29425 was more susceptible to UVC at 268.5 nm wavelength compared to other test strains (Table 1, Tables S1-S2). In general, the 500 mL bottle obtained slightly better disinfection performances compared to the 650 mL bottle.

**Table 1:**
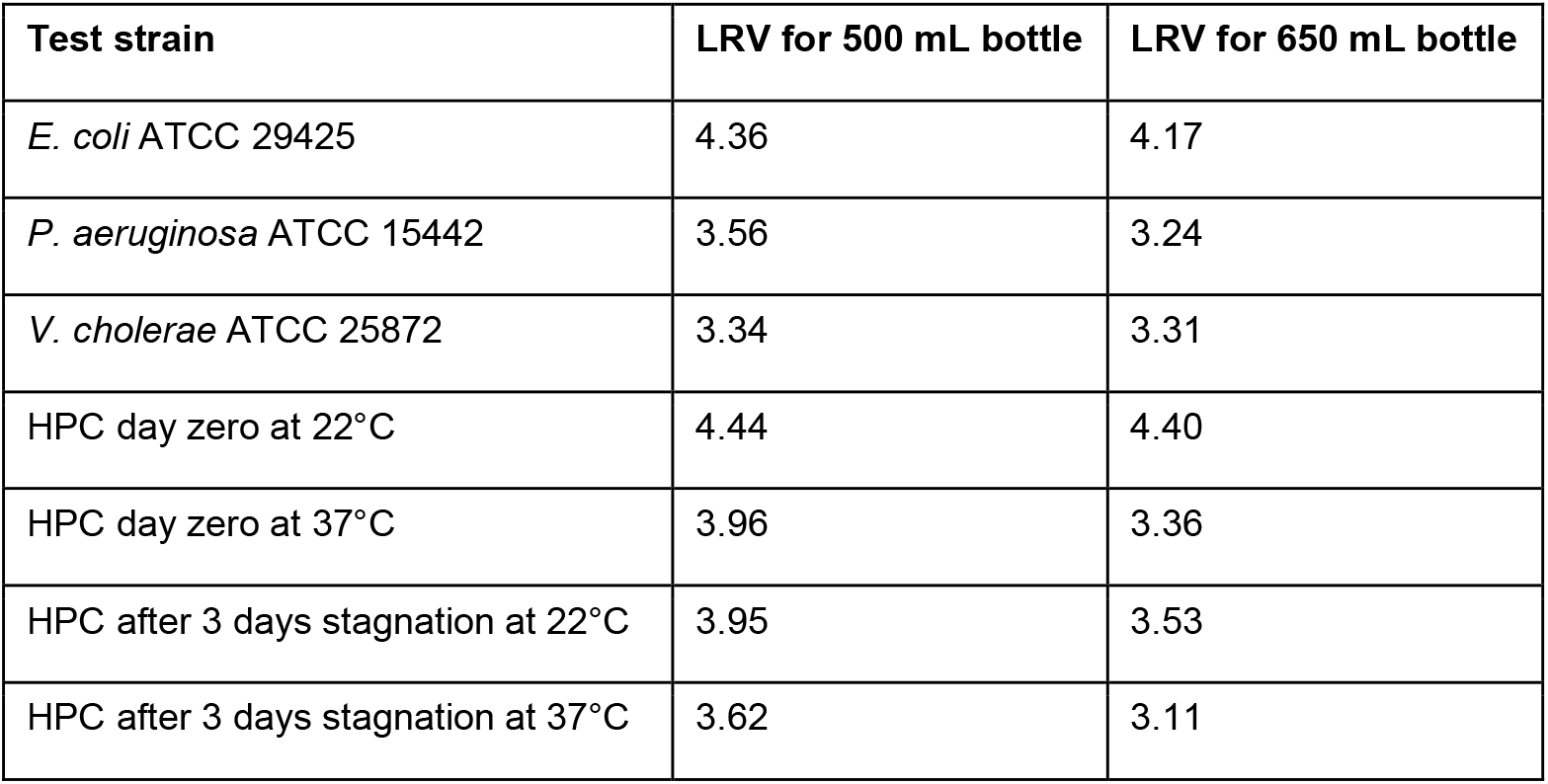
purgaty^®^ One bottles disinfection performances against bacterial strains and environmental contaminants

| Test strain | LRV for 500 mL bottle | LRV for 650 mL bottle |
| --- | --- | --- |
| *E. coli* ATCC 29425 | 4.36 | 4.17 |
| *P. aeruginosa* ATCC 15442 | 3.56 | 3.24 |
| *V. cholerae* ATCC 25872 | 3.34 | 3.31 |
| HPC day zero at 22°C | 4.44 | 4.40 |
| HPC day zero at 37°C | 3.96 | 3.36 |
| HPC after 3 days stagnation at 22°C | 3.95 | 3.53 |
| HPC after 3 days stagnation at 37°C | 3.62 | 3.11 |

**Table 2:** Identification of the HPC bacterial isolated from this study using 16S rRNA gene

| Strain name | Colony Characteristics | Accession | Organism | Closet relative | % similarity | Phylum |
| --- | --- | --- | --- | --- | --- | --- |
| Poff1 | White | MW167646 | *Acinetobacter johnsonii* | *A. johnsonii* ATCC 17909 | 99.18 | Proteobacteria |
| Poff2 | Yellowish | MW167647 | *Methylibium petroleiphilum* | *M. petroleiphilum* PM1 | 99.86 | Proteobacteria |
| Poff3 | White | MW167648 | *Acinetobacter johnsonii* | *A. johnsonii* ATCC 17909 | 99.39 | Proteobacteria |
| Pon1 | Pink | MW167649 | *Methylobacterium populi* | *M. populi* BJ001 | 99.65 | Proteobacteria |
| Pon2 | Yellowish | MW167650 | *Sphingomonas ursincola* | *S. ursincola* DSM 9006 | 99.72 | Proteobacteria |
| Pon3 | White | MW167651 | *Brevundimonas nasdae* | *B. nasdae* W1-2B | 99.28 | Proteobacteria |
| Pon4 | White | MW167652 | *Bradyrhizobium yuanmingense* | *B. yuanmingense* NBRC 100594 | 99.86 | Proteobacteria |
| Pon5 | Pink | MW167653 | *Brevibacillus choshinensis* | *B. choshinensis* NBRC 15518 | 99.45 | Firmicutes |
| Pon6 | White | MW167654 | *Brevibacillus nitrificans* | *B. nitrificans* DA2 | 99.39 | Firmicutes |
| Pon7 | Pink | MW167655 | *Roseomonas mucosa* | *R. mucosa* MDA5527 | 99.72 | Proteobacteria |
| Pon8 | White | MW167656 | *Brevibacillus nitrificans* | *B. nitrificans* DA2 | 99.25 | Firmicutes |
| Pon9 | Pink | MW167657 | *Methylobacterium rhodesianum* | *M. rhodesianum* NCIMB 13779 | 99.23 | Proteobacteria |
| Pon10 | White | MW167658 | *Cupriavidus lacunae* | *C. lacunae* S23 | 98.84 | Proteobacteria |
| Pon11 | White | MW167659 | *Caulobacter segnis* | *C. segnis* ATCC 21756 | 99.43 | Proteobacteria |

**Note:** Poff: purgaty study isolates obtained from UVC off conditions (No UVC applied). These were the most abundant.
Pon: purgaty isolates obtained from UVC on conditions (UVC applied)

### 3.2 Disinfection Performance Against HPC Bacteria

HPC bacteria were present in all untreated water samples with concentrations ranging from 5.0 x10^5^ to 5.67 x 10^5^ CFU/mL (Table S3). Disinfection using the purgaty^®^ One system reduced the microbial load of heterotrophic bacteria by > LRV3 (99.9% reduction) (Table 1, Table S3). Additionally, there was HPC bacterial selection by UVC exposure. The selection of UVC resistant bacteria is not new, as some strains obtained in this study such as Methylobacterium have been isolated previously [20].

#### 3.2.1 Identification and Phylogenetic Analysis of HPC Bacteria

Eleven bacterial monocultures with distinct characteristics isolated from UVC on condition were selected for molecular identification. They were identified as: *Methylorubrum populi*, *Sphingomonas ursincola*, *Brevundimonas nasdae*, *Bradyrhizobium yuanmingense*, *Brevibacillus choshinensis*, *Brevibacillus nitrificans*, *Roseomonas mucosa*, *Methylorubrum rhodesianum*, *Cupriavidus lacunae* and *Caulobacter segnis* (Table 2). Two most abundant isolates were however obtained under non-UVC treated conditions (UVC not applied)). These were identified as *Acinetobacter johnsonii* and *Methylibium petroleiphilum*. These two strains were UVC sensitive, thus accounting for the high HPC bacteria decontamination (Table 1, Table S3). All Isolates belonged to either Phylum Proteobacteria or Firmicutes (Table 2).

Taxonomic classification of HPC isolates using 16S rRNA gene identified *Acinetobacter johnsonii* and *Methylibium petroleiphilum* to be most dominant in untreated water. These two representatives of Phylum Proteobacteria were sensitive to UVC irradiation and thus not selected by UVC (+UVC condition). Strain Poff1, identified as *Acinetobacter johnsonii* which was isolated under UVC off condition (-UVC) belongs to the same Genus as *Acinetobacter baumannii*, a multidrug resistant nosocomial pathogen of global concern [21]. The study sought to confirm if the two species did not cluster together to rule out any concerns regarding the presence of *Acinetobacter johnsonii* in drinking water. Phylogenetic analysis revealed that they do not cluster together (Fig. S1). Further, based on literature from previous studies, *A. johnsonii* has been confirmed to rarely causes human infections and has been found to be sensitive to virtually all antibiotics [22].

## 4. Discussion

Data from this study revealed >LRV3 in test bacteria, further supporting accumulating evidence for high disinfection activities of portable UVC devices [14]. The study demonstrated high disinfection performance against *E. coli* ATCC 29425 (99.99% reduction). This can be attributed to relatively low %GC content of *E. coli* (50.68%) [23] and its peak UVC sensitivity [24]. Although the Vibrio Genus has low %GC content (~47%), the presence of more elaborate DNA repair/protection processes make them less susceptible against UVC irradiation [25], thus obtaining 99.9% disinfection under similar test conditions (Table 1). This is in comparison with *P. aeruginosa* ATCC 15442 which has higher %GC content 66.17% (https://genomes.atcc.org/genomes/371a35eda24d4ddd).

Even after 3 days of water stagnation, the use of purgaty^®^ one system effectively decontaminated that HPC bacteria, obtaining <500 CFU/mL, as recommended [26]. All the UVC selected bacteria are of rare to no public health concern. For instance, *Brevibacillus nitrificans* is a heterotrophic nitrifying bacterium [27] whose genera, Brevibacillus is one of the most widespread and is found in diverse environmental habitats, including drinking water [28]. The study revealed that phylum proteobacteria as most frequent of the identified isolates (Table 1). Although drinking water has complex microbiota, previous characterization studies have confirmed that phylum Proteobacteria is the most frequent in drinking water [29].

Results from this study indicate that UVC exposure of static water in devices such as the purgaty^®^ One systems (cap+bottle) can reduce elevated initial bacterial loads. These devices could be useful in environments where people are vulnerable to pathogens. Applications include but are not limited to those related to travel medicine, healthcare facilities where patients are vulnerable to opportunistic pathogens, hiking, remote military installations and regions having water potability challenges.

## 5. Conclusion

This study investigated the efficacy of the purgaty^®^ One system (bottle + cap) on bacterial inactivation in a stagnant water disinfection setup. The two types of stainless-steel water bottles (650mL and 500 mL) achieved more than 99.99% inactivation efficacy against *E. coli* ATCC 29425 after a single treatment cycle of 55 seconds preset by the purgaty^®^ brain (cap), including a Klaran UVC LED with peak wavelength of 268.5nm. For *P. aeruginosa* ATCC 15442 and *Vibrio cholerae* ATCC 25872, an inactivation efficacy of more than 99.9% was achieved. The bottles were also able to inactivate heterotrophic contaminants with more than 99.9% reduction, even after 3 days of water stagnation in the bottles. These results demonstrate the ability of consistent disinfection performances of a mobile, simple-to-use and safe consumer water bottle appliance with a UVC disinfection feature including a single UVC LED only, which has extended application potential during emergency preparedness such as for flooding situations, outdoor activities like mountain climbing, military use especially during operations in remote areas as well as consumer home use when in doubt of water potability. Lastly, these results offer fundamental evidence on how travel medicine can benefit from the use of personal UVC devices to ensure eradication of enteric pathogens.

## Abbreviations

ATCC: The American Type Culture Collection
CFU: Colony Forming Units
LRV: Log Reduction Value
PBS: Phosphate Buffered Solution
PCR: Polymerase Chain Reaction
POE: Point of Entry
POU: Point of Use
rRNA: Ribosomal RNA
TDS: Total Dissolved Solids
UVT: Ultraviolet Transmittance
UVC: Ultraviolet Subtype C

## Data availability

The 16S rRNA gene sequences obtained from this study are available through GenBank (https://www.ncbi.nlm.nih.gov/genbank/) under the accession numbers MW1676**46**-MW1676**59**. The Spectral data is available via https://doi.org/10.6084/m9.figshare.13144538.v1.

## Acknowledgements

Authors wish to thank Michelle Lottridge, Chris Scully, Amy Miller, Jeremy Abel, Stephan Gadhof, Paul Bilton, Dr. Kevin Kahn and James Peterson for providing help during research and proofreading the manuscript.

