## Supplemental materials for "Disinfection performance of a drinking water bottle system with a UVC LED cap against waterborne pathogens and heterotrophic contaminants"

**Supplementary Figures**

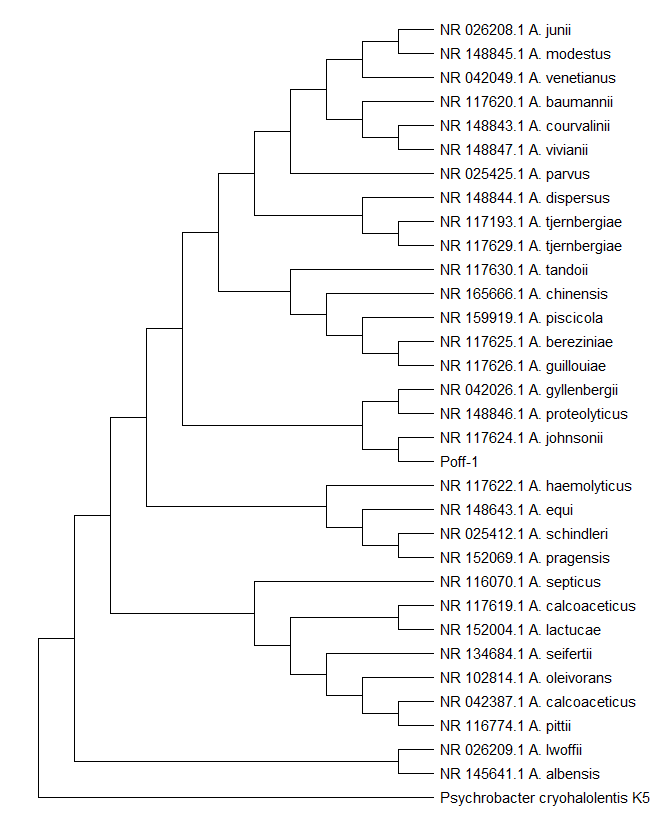

Known to be pathogenic

Known to be pathogenic

**Figure S1:** Phylogenetic analysis of *Acinetobacter johnsonii* Poff-1 isolate. Strain Poff1 did not cluster together with strains known to be nosocomial pathogens within the Genus *Acinetobacter.*

**Supplementary Tables**

**Table S1:** Disinfection performance against *E. coli* ATCC 29425 and *P. aeruginosa* ATCC 15442 obtained LRV4 and LRV3 respectively. Test water was within NSF/ANSI 55 drinking water standard and incubation was done at 37°C.

|  | | | | | Test water physicochemical characteristics | | | |
| --- | --- | --- | --- | --- | --- | --- | --- | --- |
| Bottle type | Control CFU/mL | UV on CFU/mL | LRV | % Reduction | Parameter | *E. coli*  test water | *P. aeruginosa* test water | NSF/ANSI 55  Standard |
| 500mL *E. coli* 29425 | 2.18 x 10^6^ | 9.5 x 10^1^ | 4.36 | 99.996 | UVT (%) | 96 | 96 | 98±2 |
| 650mL *E. coli* 29425 | 2.19 x 10^6^ | 1.47 x 10^2^ | 4.17 | 99.993 | pH | 7.99 | 7.98 | 7.5±0.5 |
| 500mL *P. aeruginosa* 15442 | 5.03 x 10^5^ | 1.39 x 10^2^ | 3.56 | 99.972 | Temp. (°C) | 20.4 | 20.2 | 20±2.4 |
| 650mL *P. aeruginosa* 15442 | 5.33 x 10^5^ | 3.07 x 10^2^ | 3.24 | 99.942 | TDS | 340 | 292 | 200-500 mg/mL |
|  |  |  |  |  | Turbidity | 0 | 0 | <1 NTU |
|  |  |  |  |  | Chlorine | 0 | 0 | 0 |

**Table S2:** Disinfection performance against *Vibrio cholerae* 25872 obtained LRV3 for both bottle types. Test water was within NSF/ANSI 55 drinking water standard and incubation was at 37°C.

|  | | | | | Test water physicochemical characteristics | | |
| --- | --- | --- | --- | --- | --- | --- | --- |
| Bottle type | Control CFU/mL | UV on CFU/mL | LRV | % Reduction | Parameter | Test water | NSF/ANSI 55  Standard |
| 500mL | 1.30E+05 | 5.97E+01 | 3.34 | 99.954 | UVT (%) | 96 | 98±2 |
| 650 mL | 1.30E+05 | 6.37E+01 | 3.31 | 99.951 | pH | 7.94 | 7.5±0.5 |
|  |  |  |  |  | Temp. (°C) | 18.8 | 20±2.4 |
|  |  |  |  |  | TDS | 340 | 200-500 mg/mL |
|  |  |  |  |  | Turbidity | 0 | <1 NTU |
|  |  |  |  |  | Chlorine | 0 | 0 |

**Table S3:** Disinfection performance of bottles against heterotrophic plate count bacteria at Day 0. Both bottle types obtained LRV4 at 22°C and LRV3 at 37°C. Test water was within NSF/ANSI 55 drinking water standard

|  | | | | | Test water physicochemical characteristics | | |
| --- | --- | --- | --- | --- | --- | --- | --- |
| Bottle type | Control CFU/mL | UV on CFU/mL | LRV | % Reduction | Parameter | Test water | NSF/ANSI 55  Standard |
| 500mL at 37°C | 5.25 x 10^5^ | 5.8 x 10^1^ | 3.96 | 99.989 | UVT (%) | 96 | 98±2 |
| 500mL at 22°C | 5.00 x 10^5^ | 1.8 x 10^1^ | 4.44 | 99.996 | pH | 7.92 | 7.5±0.5 |
| 650mL at 37°C | 5.67 x 10^5^ | 2.45 x 10^2^ | 3.36 | 99.957 | Temp. (°C) | 21.4 | 20±2.4 |
| 650mL at 22°C | 5.33 x 10^5^ | 2.13 x 10^1^ | 4.40 | 99.996 | TDS | 319 | 200-500 mg/mL |
|  |  |  |  |  | Turbidity | 0 | <1 NTU |
|  |  |  |  |  | Chlorine | 0 | 0 |

**Table S4:** Disinfection performance of against HPC bacteria after 72 hr of water stagnation obtained more than LRV3. The increase in TDS and subsequent decrease in UVT after stagnation led to some decrease in performance compared to LRV at Day 0.

|  | | | | | Test water physicochemical characteristics | | | |
| --- | --- | --- | --- | --- | --- | --- | --- | --- |
| Bottle type | Control CFU/mL | UV on CFU/mL | LRV | % Reduction | Parameter | Day zero | After 72 hrs. | NSF/ANSI 55  Standard |
| 500mL at 37°C | 5.27 x 10^5^ | 1.28 x 10^2^ | 3.62 | 99.976 | UVT (%) | 98 | 96 | 98±2 |
| 500mL at 22°C | 4.57 x 10^6^ | 5.17 x 10^2^ | 3.95 | 99.989 | pH | 7.98 | 8.5 | 7.5±0.5 |
| 650mL at 37°C | 5.40 x 10^5^ | 4.20 x 10^2^ | 3.11 | 99.922 | Temp. (°C) | 20.2 | 19 | 20±2.4 |
| 650mL at 22°C | 5.53 x 10^5^ | 1.63 x 10^2^ | 3.53 | 99.971 | TDS | 292 | 333 | 200-500 mg/mL |
|  |  |  |  |  | Turbidity | 0 | 0 | <1 NTU |
|  |  |  |  |  | Chlorine | 0 | 0 | 0 |
